# Monitoring mouse brain perfusion with hybrid magnetic resonance optoacoustic tomography

**DOI:** 10.1101/2022.11.23.517761

**Authors:** Wuwei Ren, Xosé Luís Deán-Ben, Zhiva Skachokova, Mark-Aurel Augath, Ruiqing Ni, Zhenyue Chen, Daniel Razansky

## Abstract

Progress in brain research critically depends on the development of next-generation multi-modal imaging tools that are capable of capturing transient functional events and multiplexed contrasts noninvasively and concurrently. A number of outstanding questions, such as those pertaining to the link between blood-oxygen-level-dependent (BOLD) signaling, oxygen saturation and underlying neural activity, could potentially be addressed by truly integrating several complementary neuroimaging readouts into one hybrid system, thus enabling a holistic view of dynamic events *in vivo*. Here we developed a hybrid magnetic resonance and optoacoustic tomography (MROT) system for murine brain imaging by incorporating an MR-compatible spherical matrix array transducer and fiber-based light illumination into a 9.4T small animal scanner, whilst further designing an optimized radiofrequency coil for whole-brain interrogation. The utility of the system is demonstrated by acquiring complementary angiographic and soft tissue anatomical contrast along with simultaneous dual-modality visualization of contrast agent dynamics *in vivo*.

## I. INTRODUCTION

Progress in understanding and disentangling the complex mechanisms of brain activity and disease progression critically depends on the development of next-generation imaging tools capable of capturing the transient functional events and multiple image contrast noninvasively and simultaneously [1–3]. Such capabilities may shed light on the link between blood-oxygen-level-dependent (BOLD) signals and oxygen metabolism in the brain and help elucidating the mechanisms of neurovascular coupling [4]. It has been long recognized that more comprehensive information can be obtained via a multi-modal imaging approach synergistically combining two or more methods [5]. For example, positron emission tomography (PET) and X-ray computed tomography (CT) are commonly combined to maximize their clinical value in oncology and other medical fields [6, 7]. Likewise, diffuse optical imaging methods can render valuable functional and molecular information in deep tissues [8], but lack the anatomical contrast and spatial resolution required for rendering quantitative information, which can be ameliorated via a multi-modal approach [9]. Multi-modal imaging is also often employed to validate the readings provided by newly developed imaging methods. Presently, approaches that can provide concurrent multimodal visualization of rapid biological dynamics, such as brain activity or perfusion, are lacking.

Optoacoustic tomography (OAT) has recently evolved as a highly versatile technique for retrieving diverse anatomical, functional and molecular information from living tissues. It provides a powerful combination between high resolution and deep (centimetre-scale) penetration not attainable with conventional optical imaging approaches [10, 11]. OAT is often hybridized with ultrasound to facilitate the image interpretation, particularly in the context of potential bed-side clinical applications [12, 13]. Co-registration of images from standalone OAT and other modalities is also routinely done for validation purposes, for which dedicated algorithms have been developed [14]. To this end, fusion of images acquired by stand-alone OAT and MRI scanners has been reported by employing software-based registration algorithms [15–17] as well as hardware-assisted protocols based on a stable dual-modal imaging support and a rigorous data acquisition protocol [14, 18]. MRI is a powerful method providing excellent soft tissue contrast, which can greatly complement the primarily vascular contrast of OAT while further benefiting from the rich functional information provided by the two modalities, such as blood oxygenation level dependent (BOLD) signals, oxygen saturation readings and contrast agent bio-distribution [10, 19].

The basic feasibility of introducing the OAT imaging hardware into the MRI bore and performing sequential image acquisitions from tissue-mimicking phantoms has recently been demonstrated [20]. In the current work, we have taken the hybrid imaging concept one step further by enabling concurrent magnetic resonance optoacoustic tomography (MROT) image acquisitions from living mice under physiological conditions.

## II. Methods

### A. Hybrid magnetic resonance and optoacoustic tomography (MROT) system

The MROT system was devised to operate on a high-field 9.4T preclinical MRI scanner (BioSpec 94/30, Bruker BioSpin, Ettlingen, Germany). A customized MR-compatible 384-element spherical ultrasound array (length: 110 mm, diameter: 90 mm) was specifically-designed to fit into the MRI bore along with a custom-made mouse holder incorporating a radio frequency (RF) coil (Fig. 1a). The MRI scanner bore (diameter: 112 mm) is horizontally aligned in parallel with a gradient coil system featuring a maximum strength of 400 mT/m [20]. The individual elements of the ultrasound array (size: ~3.5 × 3.5 mm^2^, central frequency: 5 MHz, detection bandwidth: ~80%) were arranged across a hemispherical surface with an angular aperture of 130° (solid angle 1.15π). The casing of the matrix array probe was made of non-magnetic polyetheretherketone the interference from the strong magnetic field and the RF gradients. A PEEK-made cover featuring a 30 mm central aperture was attached to the front array surface (Fig. 1b) and filled with agar (1.3% w/w) to guarantee ultrasonic transmission. The animal holder was designed with a computer-aided design (CAD) software (Inventor, Autodesk Inc., CA, USA) and 3D printed with polylactic acid (PLA, Ultimaker 2+ Connect). The cylindrical support was composed of two round bases with the same diameter (90 mm) connected with plastic screws to a bridging cradle (length: 115 mm) supporting the anaesthetized animal (Figs. 1b and 1c). The front round base (thickness: 7 mm) features an aperture that matches the probe’s diameter (Fig. 1d), whereas the back base (thickness: 3 mm) has an opening for inserting the MRI readout cable. Importantly, a customized saddle-shaped RF volumetric coil was integrated into the animal cradle close to its front base (Figs. 1b and 1c). It consists of two 22 mm diameter loops made of 1 mm thick silver wire and spaced by ~20 mm from each other. The coil resonance was tuned to 400 MHz and matched to the 50Ω BNC line with variable capacitors (ATCeramics Corp., NY, USA). A curved groove made on the animal cradle allows warming up the circulating water thus maintaining a constant body temperature of the animal under anesthesia. The support also included two ear fixation screws to improve robustness during the measurement and a specially designed breathing mask that provides gas ventilation.

**Fig. 1.**
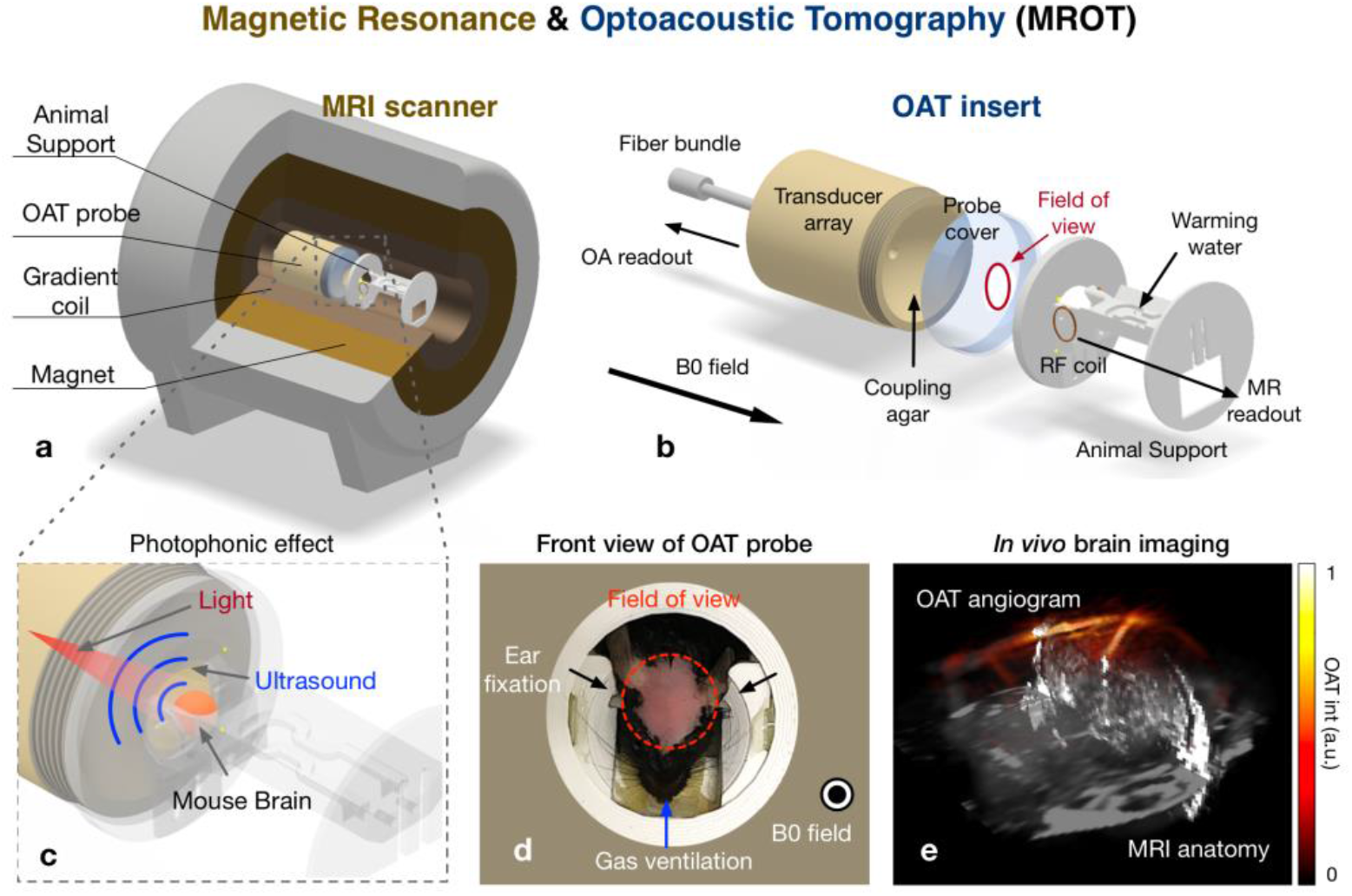
Schematic of the hybrid magnetic resonance optoacoustic tomography (MROT) system and its multimodal application for visualizing cerebrovascular perfusion in mice. (a) The optoacoustic imaging probe was inserted into a high-filed MRI scanner. (b) The insert consisted of two major components: an MR-compatible 384-element matrix array transducer and an animal support that fulfills the highly demanding *in vivo* imaging conditions. (c) Illustration of the OAT data acquisition. The mouse brain position is highlighted in red. (d) Mouse head fixation was implemented by using a pair of ear bars and specialized face mask allowing for gas ventilation. (e) The resulting multimodal image of the mouse brain shows major brain vessels visualized by OAT superimposed onto anatomical reference provided by T1-weighted MRI.

OAT imaging was performed with a tunable optical parametric oscillator (OPO)-based laser (Innolas GmbH, beam at pulse repetition frequency of 20 Hz. The laser wavelength can be tuned in the 680-900 nm range. The output laser beam was delivered to the sample by means of a customized 3 m long optical fiber bundle (Ceramoptec GmbH, Bonn, Germany) inserted into a central aperture of the array (Fig. 1b). The output ferule of the fiber bundle (length: 80 mm) was made of MR-compatible polyoxymethylene (POM), whereas the input end and the connection part were made of stainless steel and brass, respectively. The optoacoustically-generated signals detected by the matrix array transducer were simultaneously digitized with a data acquisition system (DAQ, Falkenstein Mikrosysteme GmbH, Taufkirchen, Germany) at a sampling frequency of 40 mega samples per second (MSPS) triggered with the Q-switch output of the laser. The volumetric OAT images were reconstructed from the raw sampled 512-channel data with a filtered back projection algorithm enabling real-time display during the acquisition [21]. MRI image reconstruction was performed with Paravision 6.0 (Bruker Bio-Spin, Ettlingen, Germany). All data processing steps were performed with MATLAB (R2019a, MathWorks) on a desktop computer (Intel core i7@ 2.60 GHz and RAM of 8.00 GB).

### B. Sensitivity characterization

Next, sensitivity of the MROT system for detecting standard OAT and MRI contrast agents, flowing through a 0.28 mm inner diameter tubing embedded in an agar-based phantom, was assessed (Fig. 2a). Indocyanine Green (ICG), a commonly used fluorescent probe approved for clinical use, was injected into the tubing at different concentrations (0.1, 0.25, 1, 2.5, 10, 25 and 100 *μ* M) to characterize the OAT sensitivity [22]. A GdDOTA-saline solution (Guerbet, Paris), a well-established MRI contrast agent consisting of organic acid DOTA as a chelating agent and paramagnetic gadolinium [23], was also injected into the tubing at different concentrations (0.25, 1, 2.5, 10 mM) to assess the sensitivity achieved with the customized MRI coil. All acquisitions were performed with the OAT probe positioned inside the MRI bore. MRI measurements were performed by considering a Time-of-Flight (ToF) MRI sequence featuring field of view (FOV) = 16 × 16 mm^2^, dimension of the reconstruction matrix = 100 × 100, echo time (TE) = 1.34 ms, repetition time (TR) = 10 ms, flip angle (FA) = 80 degree, slice thickness = 1 mm.

**Fig. 2.**
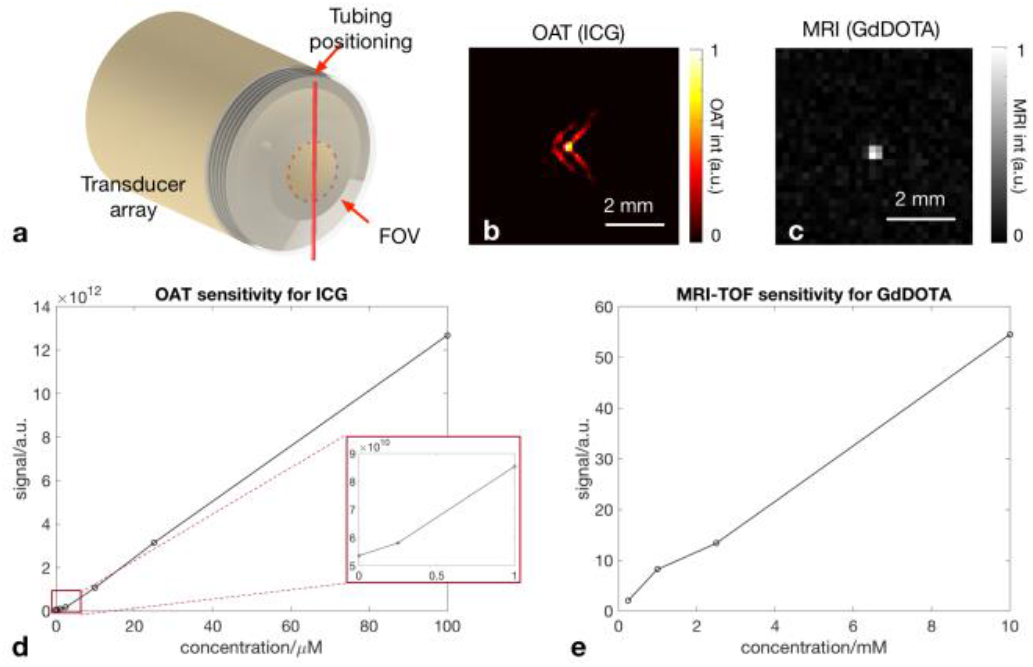
Characterization of the multimodal imaging performance. (a) Phantom experiment for characterizing the sensitivity of MROT to different concentration levels of ICG and GdDOTA solution. A tubing with 0.28 mm inner diameter (indicated in red) running the contrast agent was moved into the FOV of the spherical transducer array. (b) The tubing containing 10 μM concentration of ICG can be visualized with OAT at 800 nm excitation wavelength. (c) The corresponding ToF MRI image of 2.5 mM concentration of GdDOTA. (d) Intensity of the recorded OAT signal as a function of the ICG concentration. Minimal optoacoustic signal was detected at the concentration of 0.25 μM (indicated in an inset). (e) Similarly, the MRI sensitivity was characterized with varying GdDOTA concentrations.

### C. Animal preparation

The *in vivo* dynamic imaging performance of MROT was experimentally tested in 3-4 months old female C57BL6/J mice (n = 5, Janvier Labs, France). The animals were housed in individually ventilated cages inside a temperature-controlled room under a 12-hour dark/light cycle. Pelleted food (3437PXL15, CARGILL) and water were provided ad-libitum. All experiments were performed in accordance with the Swiss Federal Act on Animal Protection and approved by the Cantonal Veterinary Office Zurich (license number: ZH161/18). Prior to the imaging session, the mouse head was shaved to ensure undisturbed propagation of light and sound. The mice were then anesthetized via i.p. injection of a Ketamine (100 mg/kg) and Xylazine (10 mg/kg) mixture solution. Maintenance anesthesia consisting of a mixture of Ketamine (25 mg/kg) and Xylazine (1.25 mg/kg) was further injected i.p. every 45 min as described in [24]. The nose of the mice was inserted into a breathing gas mask providing a constant air flow (20%-80% oxygen-air mixture) to ensure proper breathing. Animal’s body temperature was constantly monitored with a rectal probe thermometer and kept at 36.5-37 °C throughout the experiments.

### D. In vivo hybrid MROT imaging of contrast agent perfusion dynamics

T1-weighted MRI images emphasizing anatomical structures in the mouse brain were first acquired by using a fast low angle shot (FLASH) sequence (parameters: FOV = 16 × 16 mm^2^, dimension of the reconstruction matrix = 100 × 100 × 30, TE = 3 ms, TR = 750 ms, FA = 50 degree, slice thickness = 0.5 mm). Dynamic MROT imaging was then performed during the administration (i.v.) of the ICG-GdDOTA mixture (100 *μ*1, 1.29 mM ICG and 0.5 M GdDOTA). At this stage, a single-slice ToF MRI sequence featuring a very short TR facilitated recording the blood-flow-related spin enhancement within a two-dimensional plane in the mouse brain [25] (parameters: frame rate: 1 fps, FOV = 16 × 16 mm^2^, dimension of the reconstruction matrix = 100 × 100, TE = 1.34 ms, TR = 10 ms, FA = 80 degree, slice thickness = 1 mm). Simultaneous OAT images were acquired by setting the laser wavelength to 800 nm, approximately corresponding to the peak absorption of ICG in blood, and the frame rate to 20 Hz. The reconstructed FOV for OAT images was set to 10 × 10 × 8 mm^3^, which was sufficient to cover most of the brain structures. The concurrent MROT data acquisition lasted 2 minutes and included a brief baseline recording. The mixed ICG-GdDOTA solution was injected at t = 23 s during the acquisition period. Approximately 10 minutes after the injection of the ICG-GdDOTA solution, a 3D MRI angiography map over the whole brain region was acquired with a multi-slice ToF MRI image sequence (parameters: FOV = 16 × 16 mm^2^, dimension of the reconstruction matrix = 100 × 100 × 30, TE = 1.36 ms, TR = 10 ms, FA = 80 degree, slice thickness = 0.5 mm).

### E. Hybrid MROT image registration and data analysis

Although OAT and MRI images were acquired simultaneously, the datasets were not automatically aligned in a unified common ordinate since exact position and relative orientation of the matrix array transducer with respect to the RF coil are unknown. Co-registration of the images rendered with both modalities was thus performed to facilitate accurate data analysis. For this, the MRI images were set as the fixed space and the OAT images as the moving space. The T1-weighted MRI images and the ToF MRI images were inherently coregistered since the same configuration regarding FOV, reconstruction grid and orientation was set (section II.D). Considering the fact that blood vessels constitute the major image contrast and are highly visible for both OAT and ToF MR angiograms, the vascular structures on the surface of the cortex were used as the major reference landmarks for the image registration [25]. A 3D volumetric affine transformation that maps the OAT images to the MRI images was generated by using the open-source platform BioImageSuite (www.bioimagesuit.org) [15, 25]. A single slice (2D) ToF MRI image was recorded continuously for capturing the contrast agent dynamics, which was co-registered to the equivalent slice of the volumetric OAT image (Fig. 3, right panel). Before data analysis, a spatial adjustable Wiener filter and a temporal Gaussian filter were applied to the ToF MRI for noise reduction. Subsequently, a region of interest (ROI) was identified according to the T1-weighted MRI anatomy to facilitate the analysis the ICG-GdDOTA contrast agent uptake. Post-processing steps and ROI analysis were done with MATLAB (R2019a, MathWorks) on a desktop computer (Intel core i7@ 2.60 GHz and RAM of 8.00 GB).

**Fig. 3.**
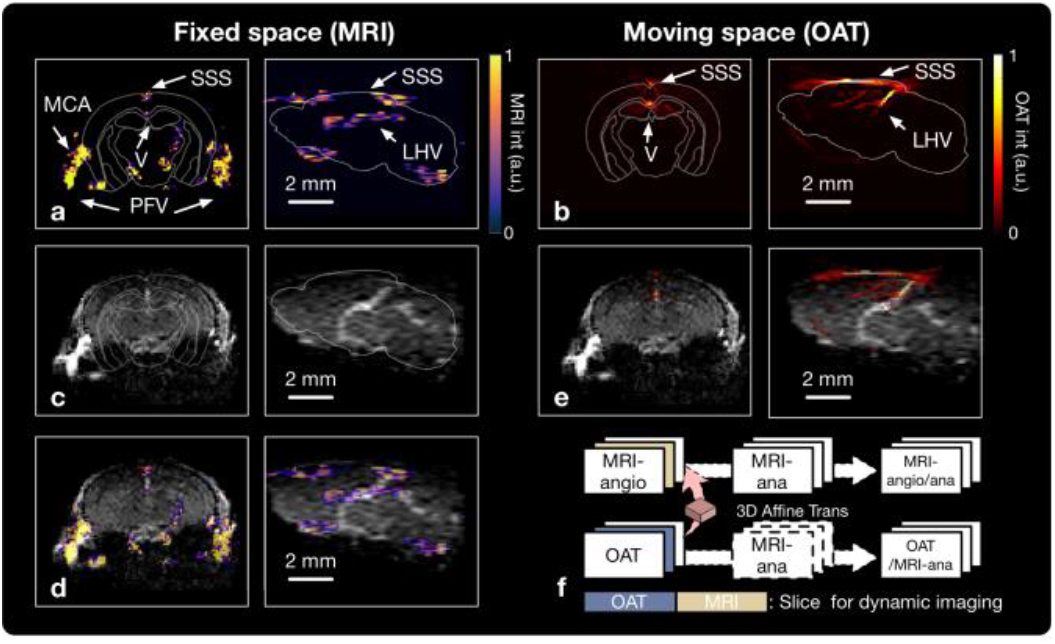
Image registration between MRI and OAT. In the left panel, both three-dimensional ToF MRI images (a) and T1-weighted images (c) are shown, representing the brain angiography and anatomy respectively. (a) and (c) were inherently co-registered, since they used the same FOV, reconstruction grid and orientation during the data acquisition step. The MRI angiography was overlaid with the T1-weighted image to better localize the signal (e). On the right panel, the OAT image shows light absorption map of the brain (b) attaining similar information as the MRI angiography. (d) overlaid OAT and T1-weighted MRI images. (f) The image co-registration protocol. Affine transformation was performed on the OAT images to map the moving image space of OAT into the fixed space given by MRI images. (SSS-Superior Sagittal Sinus, LHV-Longitudinal Hippocampal Vein, V-Ventricle, PFV-Posterior Facial Vein, MCA-Middle Cerebral Artery)

## III. Results

### A. Sensitivity characterization of MROT

First, system detection sensitivity was quantitatively characterized by using a tubing-containing phantom (Fig.2a). The cross section of the tubing filled with ICG (10 μM) and GdDOTA (2.5 mM) were visualized via OAT and MRI, respectively (Fig.2b and 2c). With the MROT insert inside the MRI bore, the measured ICG signal at 800 nm as a function of concentration when no averaging is performed is shown in Fig.3d. The minimum detectable concentration is approximately 0.25 *μ* M, similar to the corresponding OAT detection thresholds without the presence of MRI scanner [13], i.e., the interference from the magnetic field and the RF gradients appears not to have a strong effect on the OAT sensitivity. In a similar manner, the MRI signal as a function of GdDOTA concentration (in the 0.25-10 mM range) is displayed in Fig. 3e. The minimum concentration of GdDOTA that could be visualized with ToF MRI was approximately 2.5 mM, which is inferior to OAT by about three orders of magnitude.

### B. Image co-registration between the modalities

To facilitate the analysis of dynamic multimodal data, OAT images were successfully co-registered with different types of MRI images (Fig. 3). Vascular networks could be observed by both modalities. Specifically, both superior sagittal sinus (SSS) and longitudinal hippocampal vein (LHV) were clearly visualized with both ToF MRI (Fig. 3a) and OAT (Fig. 3b). Due to the high sensitivity of the surface coil, the ventricle, the posterior facial veins (PFVs) and middle cerebral artery (MCA) on both sides were further resolved in the ToF MRI images (Fig. 3a). On the other hand, excellent soft-tissue anatomic contrast was achieved with the 3D T1-weighted MRI images (Fig. 3b). After registration, both ToF MR angiograms and OAT images could be accurately overlaid over the T1-weighted MRI images (Fig. 3c and 3e), leading to a better localization of structures thus facilitating data analysis.

### C. Simultaneous dynamic MROT imaging

The kinetics of the mixed contrast agent containing ICG (1.29 mM) and GdDOTA (0.5 M) was captured by using concurrent OAT and ToF MRI recording. The time-lapse ToF MRI image slices were compared with the corresponding single slices from the four-dimensional OAT data at the same position (Figs. 4a, b, Supporting material S1, S2). Overall, signal intensity enhancement trend was observed with both modalities. However, the highest ToF MRI signal was observed in a region close to the bottom part the brain e.g. in the PFVs, while the maximum OAT signal intensity corresponds to a region around the top of the brain e.g. around the SSS. In fact, such discrepancy is not related to the differences in the contrast agent bio-distribution but rather the different distribution of the magnetic field generated by the RF coil as compared to to the excitation light fluence inside the tissue. ROI analysis was performed by selecting two regions around the SSS and Ventricle to compare the observed time courses of the detected ICG-GdDOTA signal with both modalities (Figs. 4c, d, Supporting material S1, S2). The arrow in Fig. 4d indicates the time points corresponding to the i.v. injection. After applying a Gaussian smoothing filter (sigma value = 3, length = 5), an increase in both MRI and OAT signals was observed for both selected ROIs following the contrast agent injection at t = 23s. The rising edges of both curves reasonably match. Due to different uptake performance of ICG and GdDOTA, the curves exhibit different behavior at a later stage (Fig. 4c, d). The drop of GdDOTA signal at t ≈ 55 s and t ≈ 75 s can be also explained by a slight motion of the animal head leading to the attenuated signal, a phenomenon known to significantly affect the MRI image quality [26].

**Fig. 4.**
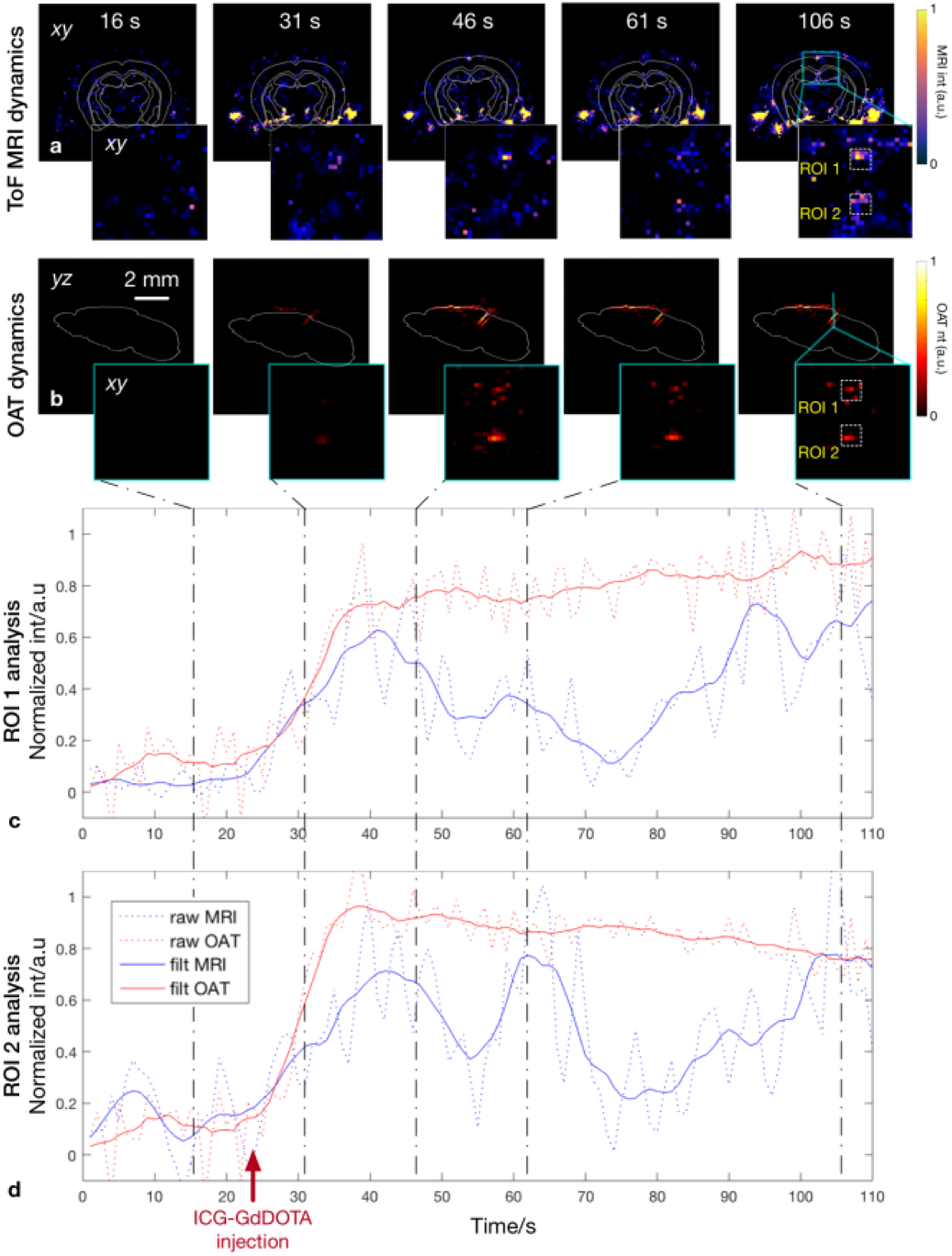
The hybrid MROT system visualizes perfusion of two contrast agents simultaneously injected at t = 23 s (indicated with a red arrow). (a) The recorded ToF MRI images sequence showing contrast enhancement by GdDOTA. (b) The corresponding time-lapse OAT images showing the increasing concentration of ICG. The dynamic ToF MRI and OAT images at 16, 31, 46, 61, 106 second time points are captured and displayed in parallel. (c, d) The time courses of OAT (red) and MRI (blue) signal intensity are plotted in two ROIs, as indicated in (a) and (b).

## IV. Discussion and conclusion

Similar to other established molecular imaging methods such as PET, the fast evolving OAT technology has entered a new era that emphasizes the need for a multimodal strategy augmented by another high-resolution anatomical imaging method featuring good soft tissue contrast. In this work, the newly developed hybrid MROT system was shown to fully retain the original advantages of standalone OAT while allowing the parallel acquisition of various MRI sequences. OAT capitalizes on the high absorption of light by blood for generating good angiographic contrast as well as capturing disease-related hemodynamic and vascular changes [27, 28]. Note that by exciting tissues at multiple optical wavelengths, the multi-spectral OAT additionally provides functional information, such as oxygen saturation readings, as well as good sensitivity for molecular imaging of targeted agents featuring distinctive absorption spectrum. OAT has also been shown to attain ultrafast imaging speeds not available with other modalities [1, 10, 20, 29]. However, due to the strong attenuation of light within living tissues, penetration depth of OAT is typically limited to a few millimeters to centimeters (Fig. 4). In contrast, MRI is a powerful tool for whole-body imaging and offers versatile tissue-specific anatomical and functional contrast mechanisms such as proton density, longitudinal relaxation time T1 and transverse relaxation time T2. Exogenous contrast agents can further enhance the MRI capabilities [23, 30]. Therefore, the combination between OAT and MRI is expected to bring synergistic advantages and complementarity in terms of spatiotemporal resolution, penetration depth, contrast, as well as multiplex anatomical, functional and molecular readouts [9].

The basic technical feasibility of integrating between OAT and MRI has previously been demonstrated in phantom experiment [20], however no *in vivo* application of the hybrid methodology has so far been implemented. In this work, we have successfully developed a hybrid MROT system that allows for *in vivo* application. This has been enabled by designing dedicated hardware components and imaging protocols. In particular, a copper shielded 384-element transducer array was developed with a large FOV covering whole-brain imaging while protecting the ultrasound detection path from strong MRI field gradients. We also customized a saddle-shaped volumetric coil which can acquire the MRI signals within a brain region of a similar size acquired by OAT. A 3D printed bridging cradle was developed, featuring compact design, stable fixation and sufficient life support for longitudinal *in vivo* measurement. The lying animal was warmed with the underlying running warm water and received sufficient gas ventilation. We then performed a concurrent dynamic MROT imaging to monitor brain perfusion after administrating the ICG-GdDOTA mixture solution. As shown in Fig.4, the time courses of both contrast agents were in a good agreement, thus further validating the functionality of the MROT system. Although integration between optical imaging methods, such as fluorescence imaging and diffuse optical tomography, and MRI has previously been reported [9, 25, 30, 31], MROT is a unprecedented multimodal combination that achieves sub-second speed, sub-millimeter spatial resolution and a large three-dimensional FOV covering the entire mouse brain with both modalities.

Our work also showcases a workflow that enables an accurate MROT data analysis. The OAT and MRI datasets are not inherently aligned in a unified common ordinate. A volumetric affine transformation was thus generated based on similarity between the angiograms acquired by MRI and OAT (Fig.3). This affine transform was afterwards performed on the OAT data to map onto both the MRI angiographic and anatomical images. Once OAT and MRI data were in a common space, the VOI analysis of each modality in a dynamic imaging mode can be performed. Another potential application of such multimodal combination is the MRI-assisted light fluence correction for quantitative OAT. Specifically, the MRI structural information was previously suggested for estimating 3D light fluence distribution, which resulted in improved OAT signal distribution corrected for strong light attenuation in deep tissue regions [14]. Light propagation modeling in a mouse brain can be more accurately predicted with a proper segmentation algorithm, which can be greatly facilitated by high-resolution anatomical maps delivered by MRI [15, 32]. In a similar manner, a dedicated light fluence correction paradigm can potentially be implemented for the suggested simultaneous MROT platform. Unlike successive acquisitions with OAT and MRI, the fully hybridized MROT platform provides unique capabilities for functional neuroimaging where concurrent recordings are required for meaningful interpretation of the brain activity. Indeed, functional MRI (fMRI) is currently a mainstay of neuroimaging, particularly for studying the mammalian brain connectivity at the whole organ level. Its BOLD signal mainly corresponds to the distribution of deoxygenated hemoglobin (paramagnetic) yet is also influenced by other hemodynamic parameters [33]. Thereby, fMRI BOLD signal can only partially unravel the complex hemodynamic changes induced by neurovascular coupling. On the other hand, multi-spectral OAT provides a unique capability to map multiple hemodynamic parameters including the biodistributions of oxygenated and deoxygenated hemoglobin [1]. The OAT-based neuroimaging, along with its intrinsic high temporal resolution, facilitates a detailed view of the whole-brain activity. The measurement of genetically encoded calcium indicators such as GCaMP further empowers OAT with the capability to directly image neuronal activity [28].

Despite being an exciting technical achievement, the MROT platform requires further optimizations. The limited space in the scanner bore challenges positioning and fixing the OAT module with respect to the MRI scanning geometry, so that the images from both modalities must be co-registered again for each individual mouse. The mouse position with its head oriented vertically and its body lying horizontally may also affect functional recordings. In future work, the geometry of the platform will be redesigned so that the mouse will be placed in a more naturally lying position facilitating long data acquisitions. In terms of MRI data quality, a signal distortion effect was observed associated to the large volume of water (agar) encapsulated inside the probe for acoustic coupling (Fig. 3b). This problem could potentially be minimized by using heavy water (deuterium oxide) for coupling as the precession frequency of the deuterium nucleus is outside the operating MRI bandwidth.

In conclusion, we have introduced a hybrid magnetic resonance optoacoustic tomography (MROT) system and demonstrated simultaneous imaging of microcirculation in the mouse brain *in vivo*. The system was initially designed to image the mouse brain but can be adapted to image other parts of the body. The comprehensive information on hemodynamic changes provided by MROT is expected to bring new insights to functional brain studies looking at neurovascular coupling mechanisms and neurodegenerative diseases.

